# Characterizing the response to face pareidolia in human category-selective visual cortex

**DOI:** 10.1101/233387

**Authors:** Susan G Wardle, Kiley Seymour, Jessica Taubert

## Abstract

The neural mechanisms underlying face and object recognition are understood to originate in ventral occipital-temporal cortex. A key feature of the functional architecture of the visual ventral pathway is its category-selectivity, yet it is unclear how category-selective regions process ambiguous visual input which violates category boundaries. One example is the spontaneous misperception of faces in inanimate objects such as the Man in the Moon, in which an object belongs to more than one category and face perception is divorced from its usual diagnostic visual features. We used fMRI to investigate the representation of illusory faces in category-selective regions. The perception of illusory faces was decodable from activation patterns in the fusiform face area (FFA) and lateral occipital complex (LOC), but not from other visual areas. Further, activity in FFA was strongly modulated by the perception of illusory faces, such that even objects with vastly different visual features were represented similarly if all images contained an illusory face. The results show that the FFA is broadly-tuned for face detection, not finely-tuned to the homogenous visual properties that typically distinguish faces from other objects. A complete understanding of high-level vision will require explanation of the mechanisms underlying natural errors of face detection.

## INTRODUCTION

Face pareidolia is the perception of illusory faces in inanimate objects. Recently, face pareidolia has been demonstrated in non-human primates (Taubert et al. 2017), evidence that it arises from a fundamental aspect of a face detection system shared across primate species. Two key features of face pareidolia are that it is typically a spontaneous and persistent phenomenon, with the object perceived simultaneously as both an illusory face and an inanimate object, and secondly that the visual attributes perceived as illusory facial features are highly variable. Face pareidolia thus offers the potential to examine how the brain processes faces in a unique situation where face perception is decoupled from the visual properties that typically define faces as a category. In the human brain, the lateral occipital complex (LOC) (Malach et al. 1995; Kourtzi and Kanwisher 2000) and fusiform face area (FFA) (Kanwisher et al. 1997) in occipital-temporal cortex respond preferentially to either objects or faces respectively. The involvement of these category-selective regions in the visual ventral stream in processing false positive faces that have a simultaneous dual face and object identity is unclear.

Several lines of evidence indicate that the response of the FFA is tightly-tuned to faces (Kanwisher and Yovel 2006). Early fMRI studies showed that the FFA responds preferentially to faces compared to other object categories, even if the faces are schematic or imagined (Kanwisher et al. 1997; O’Craven and Kanwisher 2000; Tong et al. 2000). More recently, face-related features including identity and perceptual differences have been successfully decoded from fMRI BOLD activation patterns in the FFA using multivariate pattern analysis (Nestor et al. 2011; Goesaert and de Beeck 2013; Anzellotti et al. 2014; Axelrod and Yovel 2015; Zhang et al. 2016; Guntupalli et al. 2017). Converging evidence indicates that the response of the FFA is related to face perception. Activity in the FFA correlates with face detection and identification, but not with the identification of other complex objects (e.g. cars) that require a similar level of visual expertise (Grill-Spector et al. 2004). Similarly, the FFA displays a greater response to upright than inverted faces, consistent with the robust behavioral observation of superior face recognition for upright faces (Yovel and Kanwisher 2005). Although the face-selectivity of the FFA is well-established, it is not yet known whether the spontaneous perception of a face in an exemplar from a non-preferred category (e.g. in an inanimate object) would modulate activity in the FFA.

Given its selectivity for shape and object properties (Malach et al. 1995; Kourtzi and Kanwisher 2000; 2001), it is unclear whether activity in the LOC would differentiate between inanimate objects which contain illusory faces compared to similar objects without an illusory face. Although the object-selective LOC is also responsive to faces to some degree (Yovel and Kanwisher 2004; 2005) , unlike the FFA, activity in LOC does not correlate with behavioral measures such as the face inversion effect (Yovel and Kanwisher 2005) or the degree of perceived face-likeness for simple silhouettes (Davidenko et al. 2012). Thus, while the misperception of a face in noise produces an increase in the magnitude of activity in the FFA (Summerfield et al. 2006; Liu et al. 2014) and is suggestive of a role for the FFA in perceiving illusory faces, the possible role of object-selective cortex in processing illusory faces is more difficult to predict.

To examine the representation of illusory faces in human category-selective cortex, we used functional magnetic resonance imaging (fMRI) to record patterns of blood-oxygen-level-dependent (BOLD) activation in visual brain areas while subjects viewed pictures of objects either with or without illusory faces (**Figure 2a**). We collected 56 photographs of naturally occurring examples of face pareidolia in objects such as food, accessories, and appliances. For each object with an illusory face, we found a similar image of the same category of object but without an illusory face. Importantly, this yoked image set of 56 images was matched for object content and visual features typical of the object category but did not contain any illusory faces. In total there were 8 unique image sets, each containing 14 images (**Figure 1**), which were presented in a standard fMRI blocked design (**Figure 2b,c**). In order to comprehensively characterize the response of visual ventral areas to the perception of illusory faces in inanimate objects, we applied both multivariate pattern analysis and representational similarity analysis techniques to the fMRI data. The aims were firstly to discover which category-selective areas of visual ventral cortex are sensitive to the perception of illusory faces, and secondly to determine to what extent the representation in these regions is influenced by the presence of an illusory face.

**Figure 1.**
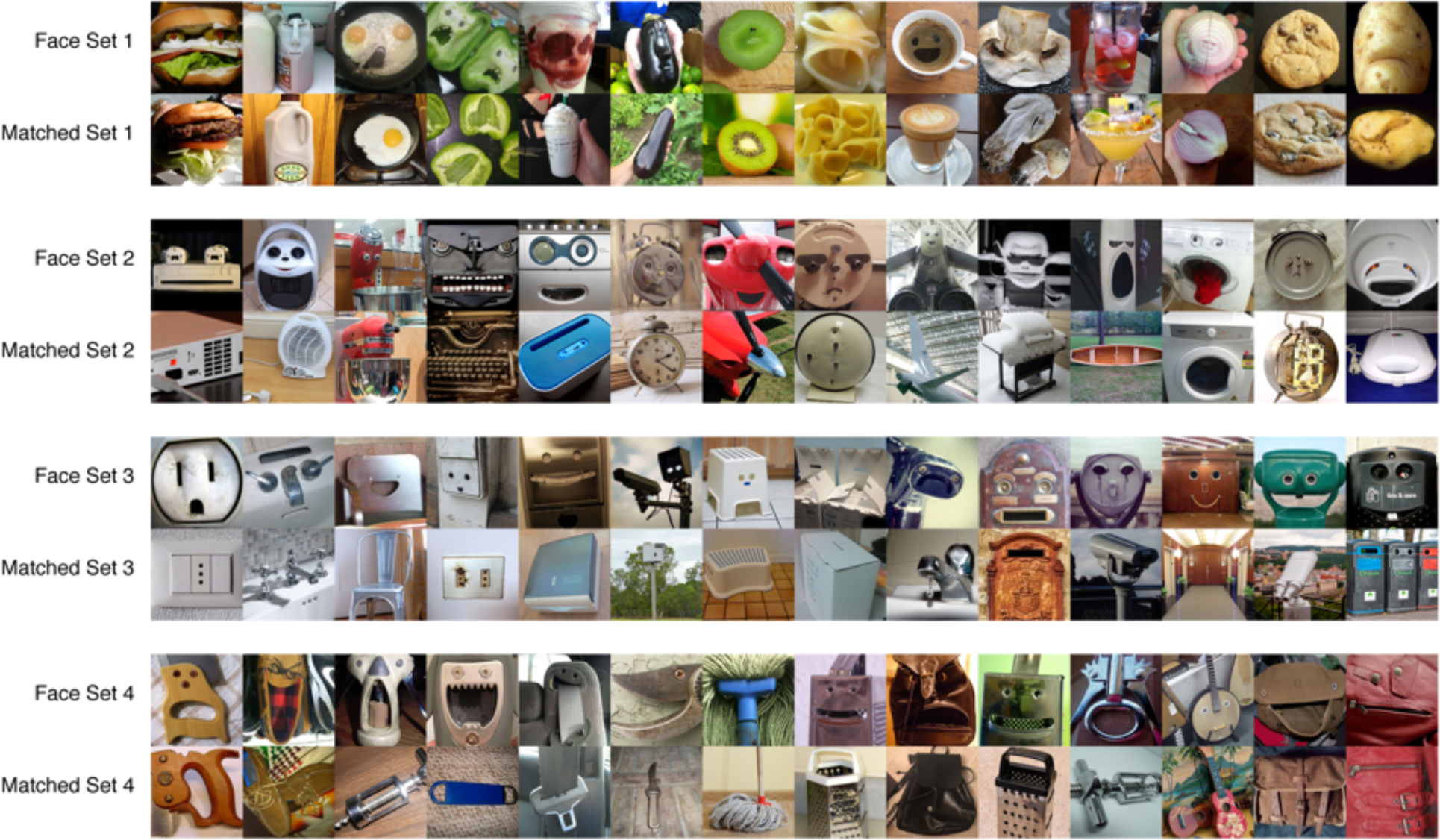
Experimental Stimuli. All 112 experimental stimuli from the 8 image sets. Four image sets contain images of illusory faces in inanimate objects, and four image sets contain images of objects matched for content and/or visual features, but without an illusory face.

**Figure 2.**
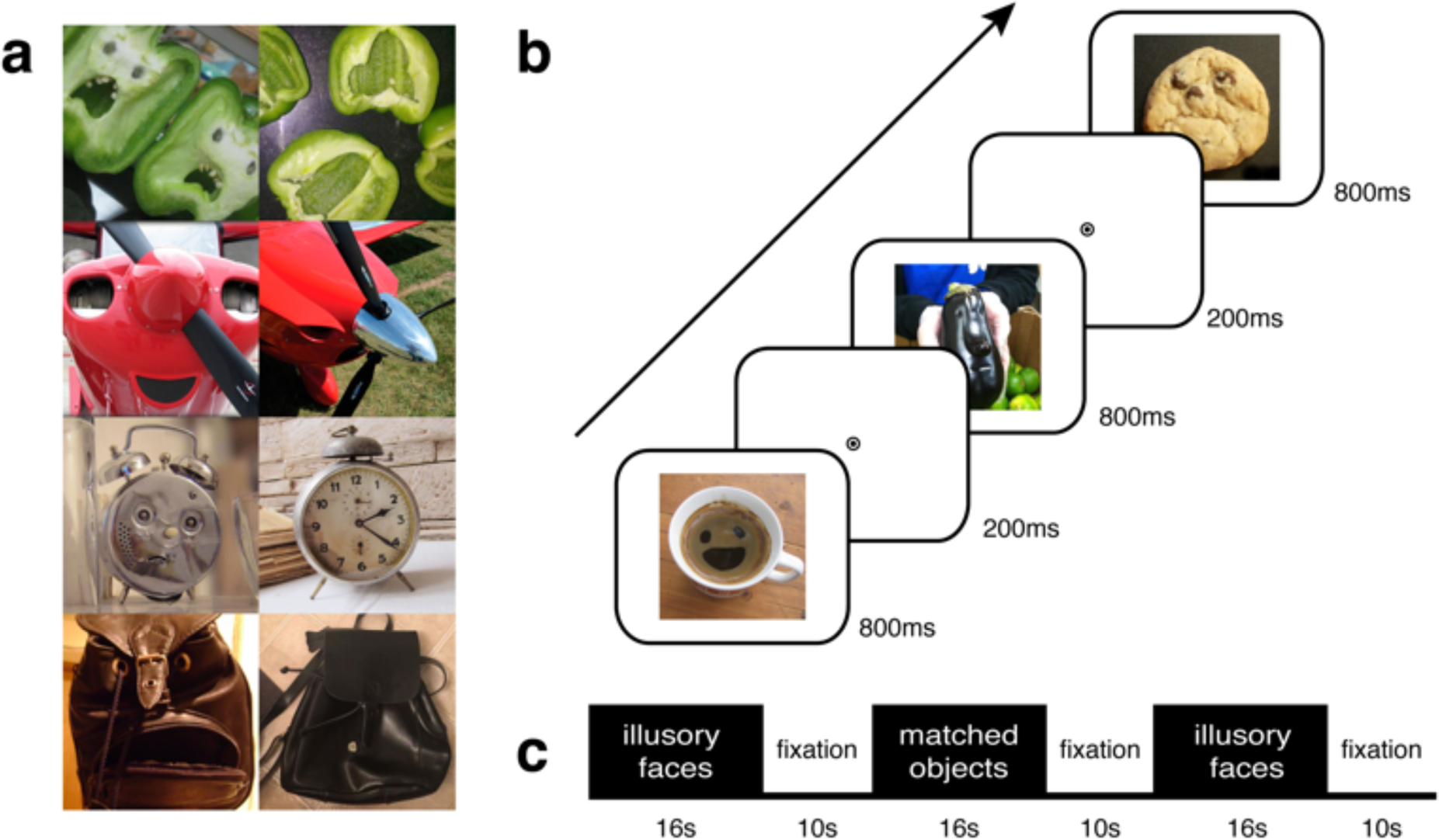
fMRI Methods. (**a**) Example stimuli: illusory faces (left) and matched objects (right). (**b**) Trial sequence within each image block. While in the scanner, participants (**N**=9) performed a 1-back task to maintain attention, pressing a key whenever an image was shown twice in a row. (**c**) Sequence of block design. Illusory face blocks alternated with matched object blocks, separated by fixation periods.

## MATERIALS AND METHODS

### Participants

Nine participants (8 naive to experimental aims (6 female), 1 author) participated in return for financial compensation and provided informed consent. One additional participant was excluded from analysis due to excessive movement in the MRI scanner (> 20mm). Ethical approval for all experimental procedures was obtained from the Macquarie University Human Research Ethics Committee (Medical Sciences).

### MRI Data acquisition

MRI data were acquired with a 3T Siemens Verio MRI scanner at Macquarie Medical Imaging, Macquarie University Hospital. A high-resolution (1 × 1 × 1 mm) T1-weighted 3D whole-brain structural MRI scan was collected for each participant at the start of the session. Functional scans were acquired with a 2D T2*-weighted EPI acquisition sequence: TR = 2s, TE = 32ms, FA = 80 deg, voxel size: 2.5 × 2.5 × 2.5mm, in plane matrix size: 102 × 102. A partial volume containing 33 slices was collected oriented approximately parallel to the base of the temporal lobe or anterior commissure-posterior commissure (AC-PC) line for each participant, ensuring coverage of both occipital and temporal lobes.

Visual stimuli were displayed using a projector for one subject (1 author) with resolution 1280 x 800 and viewing distance 150cm. Experimental stimuli (400 x 400 pixels) subtended 10.5 × 10.5°, localizer stimuli (480 × 480 pixels) 12.6 × 12.6°. For the remaining 8 naive subjects, stimuli were displayed using a flat-panel MRI-compatible 32" Cambridge Research Systems BOLDscreen with resolution 1920 × 1080 and viewing distance 112cm. Experimental stimuli subtended 7.4 × 7.4° and localizer stimuli 8.8 × 8.8°. Behavioral responses were collected using a Lumina MRI-compatible button box.

### Experimental Design and Stimuli

Experimental code was written in MATLAB with functions from the Psychtoolbox (Brainard 1997; Pelli 1997; Kleiner et al. 2007), and run on an Apple MacBook Pro with Mac OSX. The scanning session for each subject consisted of a high-resolution structural MRI scan, 2 functional localizer runs to identify category-selective regions FFA, LOC, and PPA, and 8 experimental runs. The functional localizer was run once at the start of the session following the structural scan and once at the end of the session. The eight experimental runs for each participant were collected in-between the localizer runs.

For the experimental runs, 56 color images of natural examples of illusory faces in inanimate objects were collected from the internet. For each illusory face image, we sourced a photograph of a similar object without an illusory face and cropped it to match the first image as closely as possible (**Figure 2a**). Images were cropped to the same size, but no other manipulation was made. The 112 photographs were grouped into 8 unique image sets of 14 images each, with 4 illusory face sets and 4 matched object sets (**Figure 1**). Stimuli were shown centered on a gray screen with a central black and white fixation bullseye (0.4° diameter for subject 1, 0.3° diameter for the all other subjects). During fixation blocks the fixation bullseye remained on the gray background. Each image block was 16s in duration, followed by a 10s fixation period. Within each block, every image was shown in random order for 800ms followed by a 200ms inter-stimulus-interval (**Figure 2b**). The order of presentation of the images within each block was yoked between the face image sets and their matched object sets. A new yoked random order was generated each time an image set block was repeated in the session. For each block there were 16 images: 14 unique images, and an additional 2 images which were repeated for the 1-back task. The 1-back task was yoked between illusory face and matched object blocks, such that the repeated images on a given block presentation were identical between the matched image sets. Two 1-back trials occurred during each 16s block to maintain attention, once in the first half and once in the second half. Participants pressed a key when they saw the same image twice in a row. Feedback on task performance was given at the end of each run after scanning was complete. Overall task performance was high across subjects and experimental runs (*Mean accuracy* = 98.8%, *SD* = 1.5%).

The order of the experimental blocks alternated between illusory objects and matched objects in an ABAB/BABA design (**Figure 2c**), with the starting block type counterbalanced across runs for each participant. The order of block presentation within each run was randomized within this design with the additional restrictions that (1) a matched image set never directly followed its yoked face set and vice versa, and (2) each image set was cycled through once in the run in a random order before being repeated. Each of the 8 unique image sets was repeated twice per 7-minute experimental run. Experimental runs began with a 4s fixation period.

The functional localizer stimuli were color pictures of faces, places, and objects. 54 images for each category were selected from The Center for Vital Longevity Face Database (Minear and Park 2004), the SUN397 database (Xiao et al. 2010), and the BOSS database (Brodeur et al. 2010; 2014) respectively. Scrambled objects for localizing object-selective region LOC (Kourtzi and Kanwisher 2000) were pre-generated in MATLAB by randomly scrambling each object image in an 8 x 8 grid and saving the resulting image. For each stimulus class, there were 3 unique blocks of 18 images. Every time a block was run, the images were presented in a random order and two random images were repeated twice for the 1-back task. Participants performed a 1-back task as for the experimental runs, pressing a key each time an image was repeated twice in a row (*Mean accuracy* = 96.8%, *SD* = 3.8%). Each of the 20 images within a block (18 unique + 2 repeats) was shown for 600ms followed by a 200ms inter-stimulus interval. The 16s stimulus blocks alternated with 10s fixation blocks. Each of the 4 stimulus categories was repeated three times per 5-minute localizer run, once per unique image set. The order of block presentation was in a pseudorandom order, with two different orders counterbalanced across runs for each participant. Localizer runs began with a 4s fixation period before the first stimulus block.

### fMRI data preprocessing

Minimal preprocessing of the MRI data was conducted using SPM8. For each observer, fMRI data for all experimental and localizer runs was motion corrected and co-registered to their structural scan. No normalization or spatial smoothing was applied, and all analyses were conducted in the native brain space of each subject.

### Region-of-interest ( ROI) definition

Cortical reconstruction was performed using Freesurfer 5.3 from the structural scan for each subject. V1 (*range:* 886-1244 voxels, *M*=1094) was anatomically defined in each observer based on their individual surface topology using cortical surface templates applied to their inflated cortical surface (Benson et al. 2012). The category-selective regions-of-interest (ROIs) were functionally defined using standard procedures from the independent localizer runs (Kriegeskorte et al. 2009). FFA (*range:* 29-78 voxels, *M*=58) was defined as the contiguous cluster of voxels in the fusiform gyrus produced from the contrast between faces versus objects and scenes. LOC_scr_ (*range:* 7-64 voxels, *M*=39) was defined as activation on the lateral occipital surface from the contrast between objects and scrambled objects. For comparison, LOC_ofs_ (*range:* 43-245 voxels, *M*=113) was defined as activation on the lateral occipital surface for the contrast of objects versus faces and places. Finally, PPA (*range:* 12-170 voxels, *M*=75) was defined as the peak cluster of activation in the parahippocampal gyrus produced by the contrast between scenes versus faces and objects (Kauffmann et al. 2015). Overlapping voxels were permitted between LOC_scr_ and LOC_ofs_ (range: 2-31 voxels, *M*=12), but not between the other regions.

### General linear model

To estimate the beta weights for each experimental condition, the functional MRI data from the experimental runs was entered into a GLM in SPM8 with a separate regressor for each of the 8 stimulus blocks, producing one parameter estimate (β weight) per condition per run (64 in total per participant). Fixation was not explicitly modelled but provided an implicit baseline. Percent BOLD signal change (**Figure 3a**) was calculated for each participant and region-of-interest across all face vs. matched object blocks using SPM8 and the marsbar (Brett et al. 2002) package.

**Figure 3.**
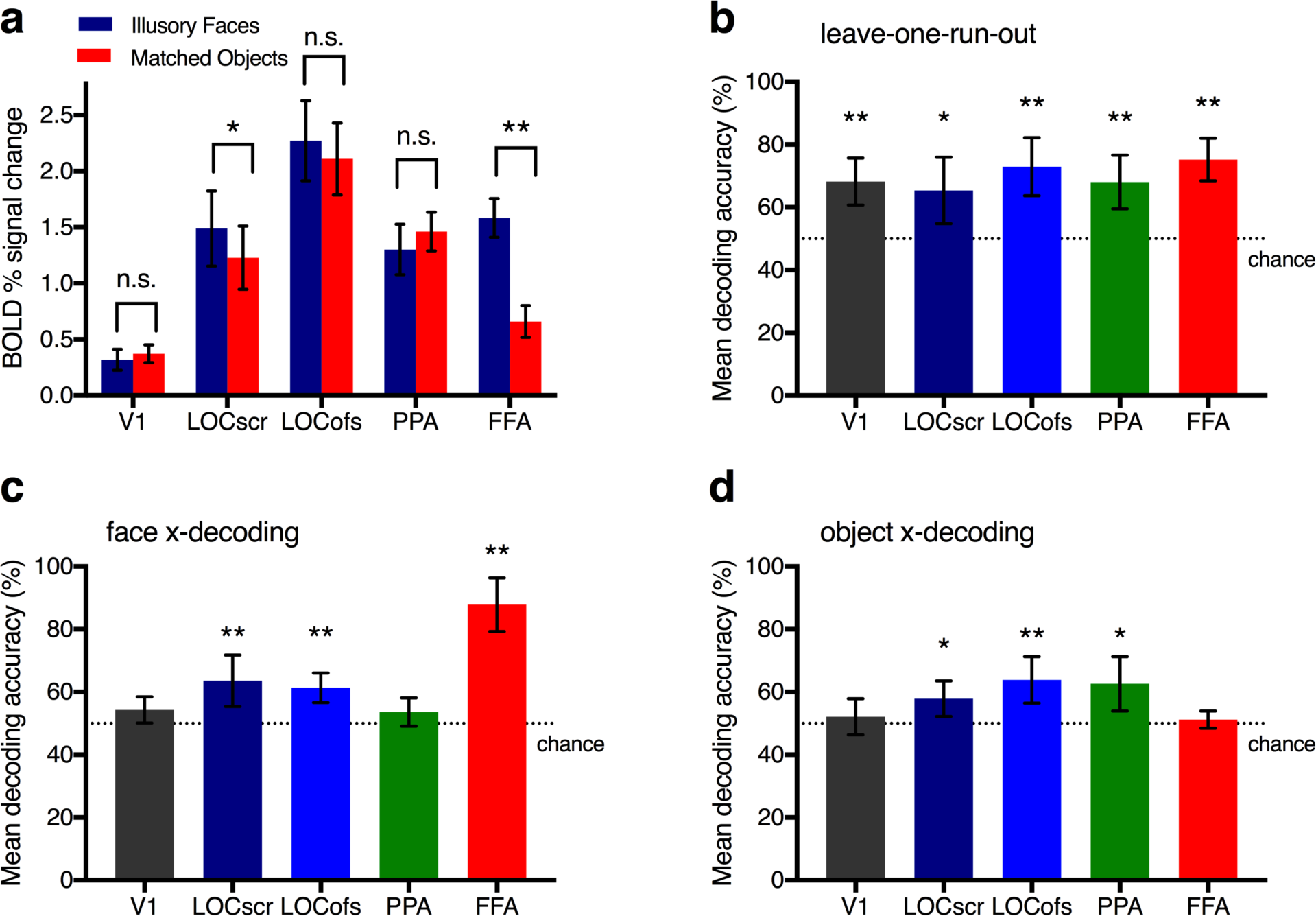
fMRI results averaged across subjects (*N*=9) for each region-of-interest. (**a**) BOLD % signal change for illusory face blocks and matched object blocks. Error bars are between-subjects *SEM*. Asterisks denote a significant difference assessed with Bonferroni paired t-tests (* α < .05, ** α < .01). **(b)** Leave-one-run-out pairwise decoding of the 8 image sets. For panels b-d, asterisks denote a significant difference from chance decoding performance (50%) assessed with Bonferroni one-sample t-tests (* α < .05, ** α < .01) and error bars are between-subjects *SD*. (**c**) Illusory face cross-decoding results. Cross-classification of illusory faces versus yoked objects without illusory faces across image sets (e.g. train classifier on Faces Set 1 vs. Matched Set 1; test classifier on Faces Set 2 vs. Matched Set 2). **( d)** Object cross-decoding results. Cross-classification of object identity across yoked image sets (e.g. train classifier on Faces Set 1 vs. Faces Set 2; test classifier on Matched Set 1 vs. Matched Set 2).

### Decoding analysis

Decoding analyses were performed separately for each participant in their native brain space in MATLAB using functions from The Decoding Toolbox (Hebart et al. 2014). Classification using a linear SVM was conducted using the beta weights for each of the 8 unique stimulus blocks. For leave-one-run-out classification (**Figure 3b**), the classifier was trained on the activation patterns for each pair of the 8 conditions for 7 runs, and tested on the one run left out of the training set. Classification accuracy was averaged across all permutations of leave-one-run-out cross-validation folds (n = 28). For face cross-decoding (**Figure 3c**), the classifier was trained on all data for a yoked stimulus set (e.g. Faces Set 1 vs. Matched Object Set 1) and tested on all data for a second yoked stimulus set (e.g. Faces Set 2 vs. Matched Objects Set 2), and then classification was repeated with the test and training sets swapped. This process was repeated for all possible yoked pairings, with classification accuracy averaged across all permutations (*n* = 6). For object cross-decoding (**Figure 3d**), two-way cross classification was conducted for each relevant pair as for face cross-decoding. For example, the classifier was trained on two different image sets either both with or without faces (e.g. Faces Set 1 vs. Faces Set 2) and tested on their yoked matched sets (e.g. Matched Objects 1 vs. Matched Objects 2), and then vice versa. Classifier accuracy was averaged across all permutations of possible pairs (*n* = 6).

### Representational similarity analysis

Representational similarity analyses (Kriegeskorte et al. 2008; Kriegeskorte and Kievit 2013) was performed in MATLAB using functions from the toolbox for representational similarity analysis (rsatoolbox) (Nili et al. 2014). Categorical models were constructed based on the stimulus design, to distinguish between the two potential outcomes of interest: whether the similarity of brain activation patterns was determined primarily by the presence of an illusory face or object content (**Figure 4c**). Blue regions indicate predicted similar patterns of activation between image set pairs, yellow regions indicate predicted dissimilar activation patterns, and grey regions indicate moderate dissimilarity. The empirical dissimilarity matrices for each brain region (**Figure 4a,b**) were calculated by 1-correlation of the BOLD activation patterns for each image set pair. Dissimilarity was scaled separately for each brain region to fall between 0 and 1 (min and max dissimilarity). Correspondence between the categorical RSA models and the dissimilarity matricies for each brain region (**Figure 4d**) were assessed by computing the correlation (Kendall tau-a) and applying condition-label randomization with FDR correction applied for multiple comparisons (*p* < .05) (Nili et al. 2014).

**Figure 4.**
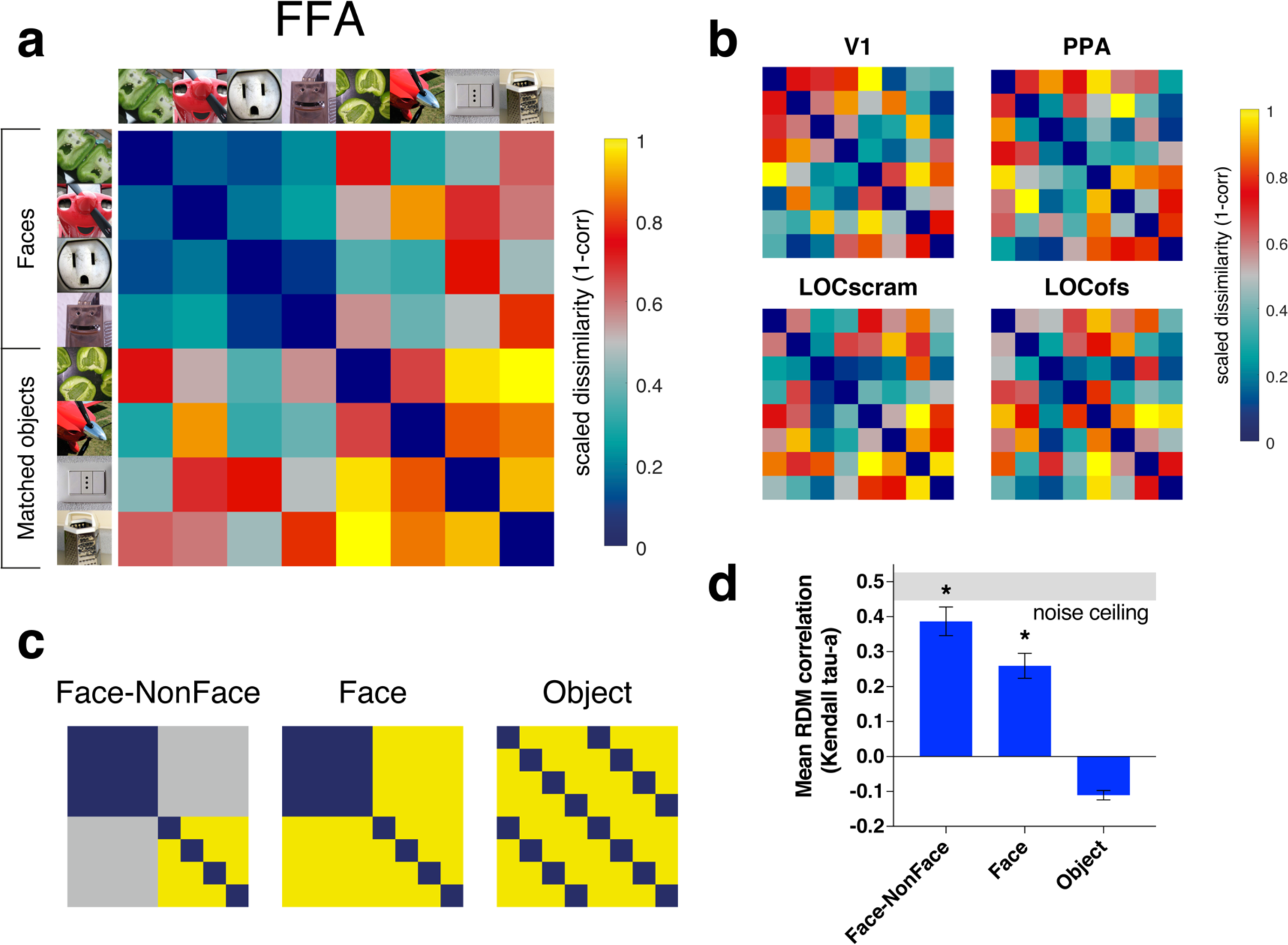
Representational similarity analysis. (**a**) Dissimilarity matrix showing the normalized pairwise dissimilarity (1 - correlation) in BOLD activation patterns evoked by each of the 8 unique image sets in the FFA. Blue indicates more similar activation patterns, yellow more dissimilar. Note that the four unique face blocks produce similar patterns of activity (upper left quadrant is blue). (**b**) Dissimilarity matrices for LOC_scr_, LOC_ofs_, PPA and V1 (legend as for panel a). (**c**) Categorical models of expected patterns of similarity in the BOLD activation patterns evoked by each image set if faces or objects dominate the underlying representation. (**d**) Correlation (Kendall tau-a) between the categorical RSA models and the FFA RDM. Mean correlation across subjects (*N*=9) is shown, asterisks denote a significant difference from chance calculated using condition-label randomization with FDR correction applied for multiple comparisons (*p* < .05). Error bars are *SEM*. The gray band represents the upper and lower boundaries of the noise ceiling, indicating the maximum possible correlation possible for any model given the noise in the data (Nili et al. 2014). Note that no correlations between the categorical models and the other regions of interest (LOC_scr_, LOC_ofs_, PPA, V1) were significant.

## RESULTS

We focused our analysis on specific predefined regions-of-interest in ventral occipital-temporal cortex. In each participant, we localized three category-selective regions: the fusiform face area (FFA), the lateral occipital complex (two versions: LOC_scr_ and LOC_ofs_), and the parahippocampal place area (PPA), using standard functional localizers (Kanwisher et al. 1997; Epstein and Kanwisher 1998; Kourtzi and Kanwisher 2000; Kravitz et al. 2011; Park et al. 2011; Kauffmann et al. 2015). As an additional comparison region, we defined V1 anatomically in each subject (Benson et al. 2012).

### Univariate analysis

To characterize the amount of activation in each region, we calculated BOLD percent signal change for all illusory face blocks versus all non-face blocks (**Figure 2a**). Statistical significance was assessed for percent signal change for illusory face versus non-face blocks on the mean activation averaged across participants for each region of interest (**Figure 2a**) using two-tailed paired t-tests with the Bonferroni correction applied for multiple comparisons (*k* = 5) to maintain α < 0.05 (α/*k* = .01). Illusory face blocks produced significantly higher activation than non-face blocks as measured by percent BOLD signal change for LOC_S_cr (*t*_(8)_= 4.25, *p* = .002814) and FFA (*t*_(8)_= 8.15, *p* = .000038). There was no significant difference in activation for illusory faces versus non-faces for LOC_ofs_ (*t*_(8)_= 2.89, *p* = .020278), PPA (*t*_(8)_= −2.68, *p* = .027728), or V1 (*t*_(8)_= −2.41, *p* = .042692).

### Multivariate pattern classification

#### Leave-one-run out classification

For an initial sanity check, we performed a standard leave-one-run-out classification analysis on the patterns of BOLD activation evoked by the 8 unique image sets. Using pairwise classification with cross-validation, a linear support vector machine was trained on the data for all runs minus one, and tested on the remaining set (Hebart et al. 2014). **Figure 2b** shows the mean classification accuracy for each region-of-interest, averaged across subjects. Decoding accuracy was assessed statistically for each ROI using two-tailed one-sample t-tests on the mean classification accuracies across participants against chance decoding performance of 50% with the Bonferroni correction applied for multiple comparisons (*k* = 5) to maintain α < 0.05 (α/k = .01). Decoding performance was significantly above chance for all five ROIs (**Figure 2b**). V1: (*t_(8)_*= 7.27, *p* = .000086), LOC_scr_ (*t_(8)_*= 4.35, *p* = .002444), LOC_ofs_ (*t_(8)_*= 7.40, *p* = .000076), PPA (*t_(8)_*= 6.33, *p* = .000226), FFA (*t_(8)_*= 11.16, *p* = .000004). Note that as the classifier is trained on examples of BOLD activation patterns evoked by each of the unique stimulus blocks using this classification method, successful classification could be based on sensitivity to low-level visual features rather than the object or face content of the image sets. It is likely that successful classification was based on different visual features in each ROI, however the consistently high classifier performance validates the high power of the fMRI design and verifies the robustness of the data for the subsequent analyses of interest.

#### Illusory face cross-classification

We used cross-decoding as the critical test for defining whether each brain region was sensitive to the presence or absence of illusory faces in inanimate objects (**Figure 2c**). A linear classifier was trained on the patterns of BOLD activation evoked by one yoked set of images (e.g. Face Set 1 vs. Matched Set 1), and tested on the brain activation patterns for a new yoked set (e.g. Face Set 2 vs. Matched Set 2). This analysis requires that the classifier be able to generalize across new images in order to correctly classify the test set, excluding the possibility of illusory face decoding based upon brain responses to the low-level visual features of particular images. Cross-decoding accuracy was assessed using two-tailed one-sample t-tests with the Bonferroni correction applied for multiple comparisons (k = 5) to maintain α < 0.05 (α/*k* = .01). Cross-classification of illusory faces was significantly above chance in LOC_scr_ (*t_(8)_*=4.97, *p* = .001096), LOC_ofs_ (*t_(8)_*= 7.21, *p* = .000092), and FFA (*t_(8)_*= 13.29, *p* = .000001), but not for V1 (*t_(8)_*=3.10, *p* = .014737) or PPA (*t_(8)_*=2.44, *p* = .040749).

#### Object cross-classification

As our stimulus set included matched objects, we also tested whether object identity could be decoded from each brain region across different photographs (**Figure 2d**). In this analysis, we ignored the presence/absence of an illusory face, and instead trained the classifier to discriminate between two different object sets that either both did or did not contain illusory faces (e.g. Face Set 1 vs. Face Set 2). The classifier was then tested on the matched sets corresponding to the two training sets (e.g. Matched Set 1 vs. Matched Set 2). Successful classification requires extrapolation based on responses to either object identity or shared visual features across different images, but it is not possible to distinguish between the two possibilities. Object cross-decoding was significantly above chance for LOC_scr_ (*t_(8)_*=4.14, *p* = .003250), LOC_ofs_ *(t_(8)_=5.63, p* = .000491), and PPA (*t_(8)_*=4.37, p = .002379), but not for V1 (*t_(8)_*= 1.12, *p* = .296317) or FFA (*t_(8)_*= 1.34, *p* = .216760). Thus both face and object cross-decoding were successful in LOC, but only faces and not objects could be cross-decoded across image sets from activity in FFA, and only objects but not faces from activity in PPA. Neither form of cross-decoding was successful in V1, yet the identity of the 8 unique image sets could be decoded from V1 activity using leave-one-run-out classification, consistent with its known sensitivity to low-level visual properties.

### Representational Similarity Analysis

To further characterize the response to illusory faces in visual ventral cortex, we applied representational similarity analysis (Kriegeskorte and Kievit 2013) to examine similarities in activation patterns for the different image sets. The representational dissimilarity matrices for each region of interest quantify the degree of similarity in the BOLD activation patterns across voxels between each pair of image sets (**Figure 4a,b**). Different models of the underlying representational space are predicted if faces or objects are the dominant organizing principle in a particular brain region (**Figure 4c**). Notably, the dissimilarity matrix for the FFA (**Figure 4a**) reveals a strong dominance of the presence of an illusory face on the brain activation patterns (upper left quadrant). Image sets containing illusory faces produce more similar responses in FFA than their matched image sets, even though the matched image sets share many more visual features with the face sets than is present between different face sets. This relationship is well-described by the face model (**Figure 4c**), which significantly correlates with the FFA dissimilarity matrix (**Figure 4d**).

Beyond a similar representation of illusory faces irrespective of their visual features, the response of the FFA is strongly modulated by the presence or absence of a face. Objects which do not have an illusory face elicit dissimilar activation patterns to each other in FFA (bottom right quadrant in **Figure 4a**), as would be expected from differences in their object identity and/or visual features. This is in sharp contrast to the activation patterns elicited by their yoked objects with illusory faces, which elicit highly similar activation patterns to each other in the FFA despite their substantial visual differences (compare upper left and bottom right quadrants in **Figure 4a**). In contrast, objects containing illusory faces are moderately dissimilar to those without illusory faces (top right quadrant in **Figure 4a**), despite their visual similarity resulting from the yoked stimulus design. Overall, the results indicate that the response of FFA is dominated by the presence or absence of a face to the extent that it overrides any influence of the similarity of visual properties or object identity in the representation.

These relationships are well-captured by the face-nonface model (**Figure 4c**), which significantly correlates with the FFA dissimilarity matrix (**Figure 4d**) and approaches the theoretical limit of the maximum possible correlation with the fMRI data as defined by the noise ceiling (Nili et al. 2014). Notably the object model fails to capture the representational structure of the visual stimuli in FFA, indicated by a lack of a significant correlation with the FFA dissimilarity matrix (**Figure 4d**). In sum, the perception of an illusory face mediates activity in FFA, regardless of the particular visual properties that compose the facial features. Although the presence of an illusory face could be successfully decoded from activation patterns in LOC_scr_ and LOC_ofs_ (**Figure 4b**), none of the dissimilarity matrices for the other regions of interest (**Figure 4b**) significantly correlated with the categorical models (**Figure 4c**), suggesting this organization of the representational space is unique to the FFA.

## DISCUSSION

We investigated the representation of illusory faces in inanimate objects throughout category-selective human ventral visual cortex. Importantly, we used a yoked fMRI design to compare patterns of BOLD activation evoked by images of objects containing illusory faces with that evoked by similar matched objects without illusory faces. We selected only naturally-occurring examples of face pareidolia in order to investigate spontaneous false positives of face detection with a high degree of variance in their specific visual properties. The cross-decoding results revealed that activity in both face (FFA) and object-selective (LOC) regions was modulated by images that elicit face pareidolia, regardless of the particular visual elements that compose the facial features and its configuration. Given that V1 is particularly sensitive to low-level visual features and is implicated in the early stages of retinotopic visual processing, it is unsurprising that the classifier failed to decode the presence of an illusory face from activation patterns in this region. We also found no evidence that activity in scene-selective PPA is mediated by the perception of illusory faces. This is notable as PPA is frequently functionally defined as voxels which exhibit a greater response to scenes than faces (Epstein and Kanwisher 1998; Kravitz et al. 2011; Park et al. 2011; Kauffmann et al. 2015), thus a reduction in activity for illusory faces might have been expected. However, there was also no difference in the magnitude of BOLD activation in PPA for illusory faces versus matched objects.

In addition to successful classification of illusory faces across image sets, activity in object-selective regions was also informative about object content. Successful object cross decoding in LOC is consistent with previous results suggesting patterns of activation in these regions are strongly associated with object identity (MacEvoy and Epstein 2009) and object shape/features (Kourtzi and Kanwisher 2000). Notably, object cross decoding was also successful in PPA, but not in FFA or V1. There is considerable influence of low-level visual features on the response of scene-selective regions (Groen et al. 2017), thus cross-classification of objects in PPA may be based upon responses evoked by shared visual features between different pictures of the same object class.

Although both FFA and LOC contained decodable information about the perception of illusory faces in inanimate objects, it is likely that the structure of the corresponding representations of these categorically ambiguous stimuli is substantially different between these two regions. Beyond their difference in preferred category (faces vs. objects) as measured by the magnitude of the BOLD response (Kanwisher et al. 1997; Kourtzi and Kanwisher 2000), there are other known functional differences. Previous studies have observed that activity in LOC does not correlate with behavioral measures of face processing (Yovel and Kanwisher 2005; Davidenko et al. 2012), while activity in FFA does (Grill-Spector et al. 2004; Yovel and Kanwisher 2005). This is consistent with our finding that the face models significantly correlated with the FFA dissimilarity matrix, and not with that for object-selective LOC. This suggests that the presence of illusory faces is a stronger predictor of the underlying representation of these objects in the FFA, while the representation in LOC is likely to involve other factors. Further, the object cross-decoding analysis revealed that activity in LOC was also informative about object identity, while object content could not be decoded from the FFA. Together these results point to genuine functional differences in the representation of illusory faces in face- and object-selective cortical regions.

The functional architecture of human ventral temporal cortex is understood to involve both clustered category-selective regions such as the FFA, and more distributed patterns of category-selectivity across larger regions of cortex (Grill-Spector and Weiner 2014). An important outstanding question is to understand the functional significance of the information the FFA contains about nonface objects (Haxby et al. 2001; O’Toole et al. 2005), which is a challenge to its face-specificity (Kanwisher 2017). A relevant point raised by our results is that the response of the FFA to inanimate objects is strongly modulated by the spontaneous perception of an illusory face. Image sets with diverse visual properties and object identities elicited similar representations if they contained illusory faces. Conversely, image sets with the same degree of variance in visual properties and object identity elicited highly *dissimilar* representations if no illusory faces were present. Notably even yoked image sets matched for visual features and object identity produced markedly dissimilar representations if only one set contained illusory faces, despite their considerable visual and semantic similarity on other dimensions. This suggests that any non-face object information contained in the FFA is secondary to the dominance of face information in its representational organization.

A significant current issue in studying the neural mechanisms of object recognition is in disentangling the degree to which category-selective responses in inferior temporal cortex reflect category representations *per se,* versus sensitivity to the low level visual features (e.g. shape, color, symmetry) that co-vary with category membership (Cox and Savoy 2003; Baldassi et al. 2013; Rice et al. 2014; Wardle and Ritchie 2014; Andrews et al. 2015; Bracci and Op de Beeck 2016; Coggan et al. 2016; Kaiser et al. 2016; Proklova et al. 2016; Bracci et al. 2017). It is intuitive that by definition, any high-level visual representations must be related to the visual features that define object categories to some degree. However, the impossibility of comprehensively accounting for all possible combinations of visual features at different levels of description entails that it is empirically challenging to provide evidence of category-representations that are not in principle reducible to some set of co-varying visual elements. The novel approach we take here is to use a stimulus class in which face perception is decoupled from the typical visual features that define real faces. Secondly, the use of a yoked experimental design means that the sets of illusory faces share more visual features with their corresponding matched objects than with each other, ensuring that the visual variance is higher within-category than across-category in our design. This is an extreme example of introducing high variance into the category exemplars (Bracci and Op de Beeck 2016; Bracci et al. 2017), as the visual features in our stimulus set are instead diagnostic of non-preferred object categories (for the FFA). A further advantage of this approach is that it does not require artificially controlling the visual content of the experimental stimuli and therefore reducing their ecological validity (Talebi and Baker 2012). Instead, in our design we exploit the high variability in naturally occurring false positives of face detection made by the primate visual system (Taubert et al. 2017).

Together our results show that the FFA strongly responds to false positive faces in inanimate objects and is tolerant to a high degree of visual variance in what constitutes a "face". Previous arguments for the face-selectivity of the FFA have been predominantly based on its preferred response to faces, typically quantified by a higher magnitude of fMRI BOLD percent signal change to faces compared to other categories (Kanwisher and Yovel 2006). In addition to human faces (Kanwisher et al. 1997), the FFA is known to respond to animal faces (Tong et al. 2000), schematic faces (Tong et al. 2000), and imagined faces (O'Craven and Kanwisher 2000). More recently, further corroborating evidence has arisen from successful decoding of face-related properties from activation patterns in the FFA such as identity (Nestor et al. 2011; Zhang et al. 2016), famous faces (Axelrod and Yovel 2015), differences in facial features and their configuration (Goesaert and de Beeck 2013), and tolerance to viewpoint rotation (Anzellotti et al. 2014). Importantly, our results show that the face-selective response in the FFA is not reducible to the specific low-level visual features that often distinguish real faces from inanimate objects. The FFA is not finely-tuned to a homogenous set of visual features. Instead, we find that the FFA is broadly-tuned to the detection of a face, independently of the particular visual features that form this perception.

Object recognition and face processing are two of the most impressive computational achievements of the human visual system, and the underlying neural mechanisms are still being revealed. To date the development of the theoretical framework for understanding the functional architecture of ventral temporal cortex and its role in high-level visual processing has focused on exemplars with distinct category membership (Kravitz et al. 2013; Grill-Spector and Weiner 2014). Our results with categorically ambiguous exemplars emphasize the complexity with which visual information is modulated by higher level perception in category-selective regions of the ventral visual stream. Further examination of natural ‘errors’ or false positives such as in the case of face pareidolia may help constrain models of high-level visual processing in the future.

## Acknowledgements

We thank Jeff McIntosh and the staff of Macquarie Medical Imaging at Macquarie University Hospital for assistance with the operation of the MRI scanner, and Robert Keys for assistance with data collection. This work was supported by an Australian NHMRC Early Career Fellowship (APP1072245) and Macquarie University Research Development Grant awarded to S.G.W.

